# Oral somatosensation shapes lingual muscle maps in macaque orofacial primary motor cortex

**DOI:** 10.64898/2026.09.10.750542

**Authors:** Shreyas Punacha, Edric D. Tsang, Andrea B. Burke, Haley Cowan, Lilah Q. Favour, Sherman Hayes, Jing-Sheng Li, Derrick Tang, Fritzie I. Arce-McShane

**Author notes:** **Corresponding author:** Fritzie I. Arce-McShane, Department of Oral Health Sciences, School of Dentistry, University of Washington, Seattle, WA, USA.

## Abstract

Oral behaviors such as chewing, swallowing, and speech depend on continuous sensory feedback from the tongue and oral tissues. Yet how oral somatosensation shapes motor cortical output to drive individual tongue muscles remains unknown. We combined electrical stimulation of 96 sites in the orofacial primary motor cortex (M1) with electromyographic (EMG) recordings from six tongue muscles in awake rhesus macaques (*Macaca mulatta*). Cortical motor maps of evoked muscle activity were evaluated under three oral sensory conditions: intact sensation, combined oral sensory nerve block, and selective preservation of tongue sensation. Orofacial M1 has a broadly distributed, strongly overlapping, and bilateral organization of individual tongue muscle representations. Under altered sensation, spatial reorganization, attenuation, and enhancement of cortical-to-muscle output emerged within a fast timescale. Selectively preserving tongue sensation neither restored the intact pattern, nor reproduced the combined nerve block pattern; instead, it generated a spatially distinct configuration of cortical-to-muscle outputs. Together, these findings demonstrate muscle-level, input-specific, and dynamic sensory shaping in orofacial M1, establishing oral somatosensory input as an active determinant of tongue-muscle output, rather than a peripheral modulator. The rapid reorganization suggests that altered sensation qualitatively reweights existing sensorimotor pathways, rather than merely scaling cortical output up or down, such that its disruption could immediately degrade tongue muscle coordination while enabling adaptation to sensory loss.

## Introduction

Essential oral functions such as mastication ^1^, swallowing ^2^, and articulated speech ^3^ require highly coordinated activation of the tongue and jaw muscles ^4–7^. This coordination requires descending motor commands from the orofacial primary motor cortex (M1) that are informed by tactile and proprioceptive information from the orofacial somatosensory cortex (S1) ^8–11^. Trigeminal afferents convey much of the sensory information from the tongue, teeth, palate, and oral mucosa ^12^. Relative to other body regions, the tongue has disproportionately large representation in the homunculus of M1 and S1, reflecting extraordinary levels of sensorimotor control required by behaviors involving the tongue.

The motor cortical representation of individual arm and hand muscles has been extensively characterized in humans and non-human primates ^13–15^. The M1 neurons project to specific motoneuron pools, with some muscle selectivity, but also a considerable overlap ^14,16,17^. Unlike the predominantly contralateral projections typical of limb motor control ^18^, M1 projections to the hypoglossal motor nuclei via the corticobulbar tracts are bilateral, with contralateral predominance ^19,20^, reflecting the need for coordinated bilateral tongue movements. Another distinction from limb muscles is that the tongue is a muscular hydrostat and not a single actuator ^21^. Four bilateral intrinsic muscles reshape the tongue (i.e., elongation, flattening, cupping) while four bilateral extrinsic muscles reposition it in space ^22^. The varied shapes the tongue assumes as it moves inside the mouth when eating or speaking involve coordination of at least eight muscles with distinct anatomical attachments and biomechanical roles, thus providing high degrees of freedom ^23^. Anatomical and physiological studies localized the macaque tongue representation in orofacial M1^19,20,24^. Suprathreshold intracortical microstimulation (ICMS) evoked non-specific tongue movements only or coordinated behaviors with the jaw such as rhythmic chewing or swallowing ^25,26^. Yet, the muscle-level organization of orofacial M1 remains undefined for the tongue. Without muscle-level mapping, we cannot fully understand how M1 controls the coordination among tongue muscles.

Studies of limb M1 show that sensory loss can alter motor cortical output over multiple timescales. Temporary ischemic nerve block rapidly increased the excitability and spatial extent of proximal motor outputs in humans ^27^. Motor cortical stimulation and behavioral training modified the effects of transient forearm deafferentation, demonstrating that neuroplastic changes remain adaptable ^28,29^. Chronic dorsal column lesions also reorganized ICMS-defined M1 maps in macaques ^30^. Similarly, lingual nerve transection altered ICMS-defined M1 tongue and jaw representations in rats ^31^, and incisor extraction changed ICMS-evoked jaw and tongue representations ^32^. In macaques, reversible inactivation of orofacial S1 and nerve block injected into sensory branches of the trigeminal nerve impaired tongue control and activity of M1 and S1 neurons ^33–35^. In humans, lingual nerve anesthesia altered tongue motor-evoked potentials ^36,37^. To date, muscle-specific mapping which can reveal how sensory input differentially weights distinct cortical-to-muscle pathways has not been done. Knowing how precisely orofacial M1 drives individual tongue muscles has important implications on the development of brain-machine interfaces and cortical stimulation therapies for rehabilitation of dysphagia or dysarthria. Muscle-level maps would enable more precise targeting, which is particularly important given the large region of M1 dedicated to the tongue.

In this study, we investigated the muscle-level organization of orofacial M1 and the graded bidirectional changes in M1-to-muscle output in awake macaques when targeted oral sensory nerves were desensitized versus when lingual sensation was spared. We hypothesized that acute sensory loss would alter muscle-specific M1 output and that selective preservation of lingual afference would produce tongue-muscle maps more similar to control than those observed during combined and targeted sensory block.

## Results

Two rhesus macaques were chronically implanted with a 96-electrode array in orofacial M1 (*S*_1_: left M1, *S*_2_: right M1) and EMG electrodes in tongue muscles bilaterally (Fig. 1b-c). To characterize muscle-level tongue representations in M1, we used subthreshold ICMS (18 *µ*A, 15 Hz) delivered through individual M1 electrodes to evoke EMG responses in genioglossus, hyoglossus, and intrinsic tongue muscles. To test whether somatosensory input shapes these tongue-muscle outputs, we conducted three recording sessions, one per day, with the order of conditions being fixed and identical across both subjects: Combined-nerve block, then Control, then Selective-nerve block (Fig. 1d-f). In combined nerve block (Fig. 1e), the lingual, inferior alveolar, buccal, and maxillary sensory nerves were all blocked, eliminating sensory input from the anterior tongue, dentition, gingiva, buccal mucosa, and palate. In selective nerve block (Fig. 1f), the lingual nerve was spared, preserving somatosensory input from the anterior tongue while silencing input from the remaining targeted oral structures. These blocks affected tactile sensation only, leaving tongue motor innervation intact. For each M1 electrode–muscle pair, we computed stimulus-triggered averages (StTAs) of EMG activity to quantify response magnitude and construct muscle-specific cortical output maps across all 96 stimulation sites.

**Figure 1.**
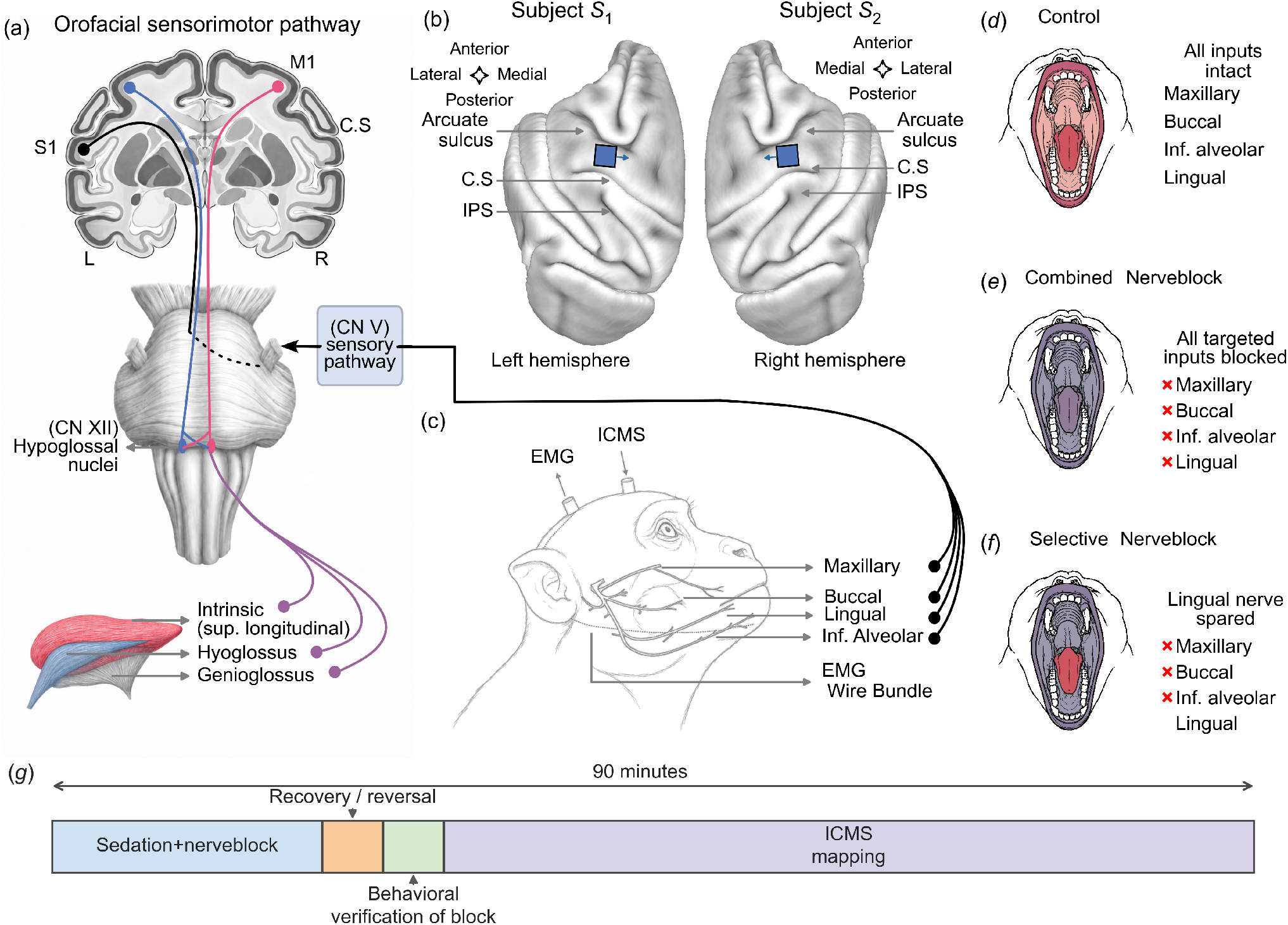
Experimental design and schema for ICMS mapping of tongue-muscle outputs during reversible oral sensory nerve block. (a) Schematic of the orofacial sensorimotor pathway: M1, S1, trigeminal (CN V) sensory, and hypoglossal motor (CN XII). Anatomical representation of tongue muscles targeted for EMG recording and oral sensory nerves targeted for nerve blocks. (b) Cortical array locations ^38,39^ in orofacial M1 of subjects *S*_1_ and *S*_2_. (c) Schematic of the trigeminal sensory branches targeted for oral nerve blocks. (d-f) Oral sensory conditions used during ICMS mapping: control, combined nerve block (Combined-NB) and selective nerve block (Selective-NB), respectively. Red and purple indicate regions with preserved and blocked tactile sensation, respectively. (g), Experimental timeline showing sedation, nerve block administration, recovery and reversal (Combined-NB and Selective-NB only), followed by behavioral verification of sensory loss and ICMS mapping within 90 min.

### Distributed and overlapping M1 tongue muscle maps

Past studies have characterized orofacial M1 sites corresponding to tongue movements induced by suprathreshold ICMS ^25,40,41^. The muscles used in the study are innervated by the hypoglossal nerve and have distinct functional roles. The genioglossus originates from the mandible and extends into the tongue. It contributes to protrusion and anterior positioning during feeding, swallowing, and speech ^42,43^. The hyoglossus extends from the hyoid into the lateral tongue and contributes to depression and retraction, including posterior tongue movement during swallowing. Intrinsic muscles originate and terminate within the tongue and comprise longitudinal, transverse, and vertical fibers. They reshape the tongue for functions such as bolus manipulation and speech articulation ^22,23^. Here, we first describe the features of muscle-level organization of orofacial M1 revealed by subthreshold ICMS under intact oral sensation (‘Control’).

Tongue movements in feeding or drinking often require bilateral activation of one or more muscles. Consistent with this, stimulation at a single electrode mostly produced responses in several tongue muscles, often bilaterally (Fig 2b-c). In *S*_1_, 86 ICMS sites (89.6%) induced StTA responses meeting the 3 SD criterion in all six muscles, while the remaining sites represented four or five muscles (Fig. 2d). In *S*_2_, 63 sites (65.6%) represented all six muscles, 24 represented five muscles, eight represented four muscles and one represented three muscles (Fig. 2e). These responses were broadly distributed across the array rather than confined to separate clusters for individual muscles; each tongue muscle was represented across a large portion of the sampled M1 region, ranging from 93.8% to 100% of sites in *S*_1_ and from 87.5% to 97.9% in *S*_2_ (Fig. 2f). The broadly distributed and overlapping tongue-muscle outputs agree with classical descriptions of intermingled face, jaw, and tongue representations in macaque orofacial M1^24,25^. Overlapping muscle representation suggests that M1 neurons connect to multiple motoneuronal pools, that may underlie mechanism for synergistic or antagonistic action of these six muscles. The evoked responses being predominantly bilateral is a sharp contrast to the contralateral responses commonly found in limb M1^44,45^.

**Figure 2.**
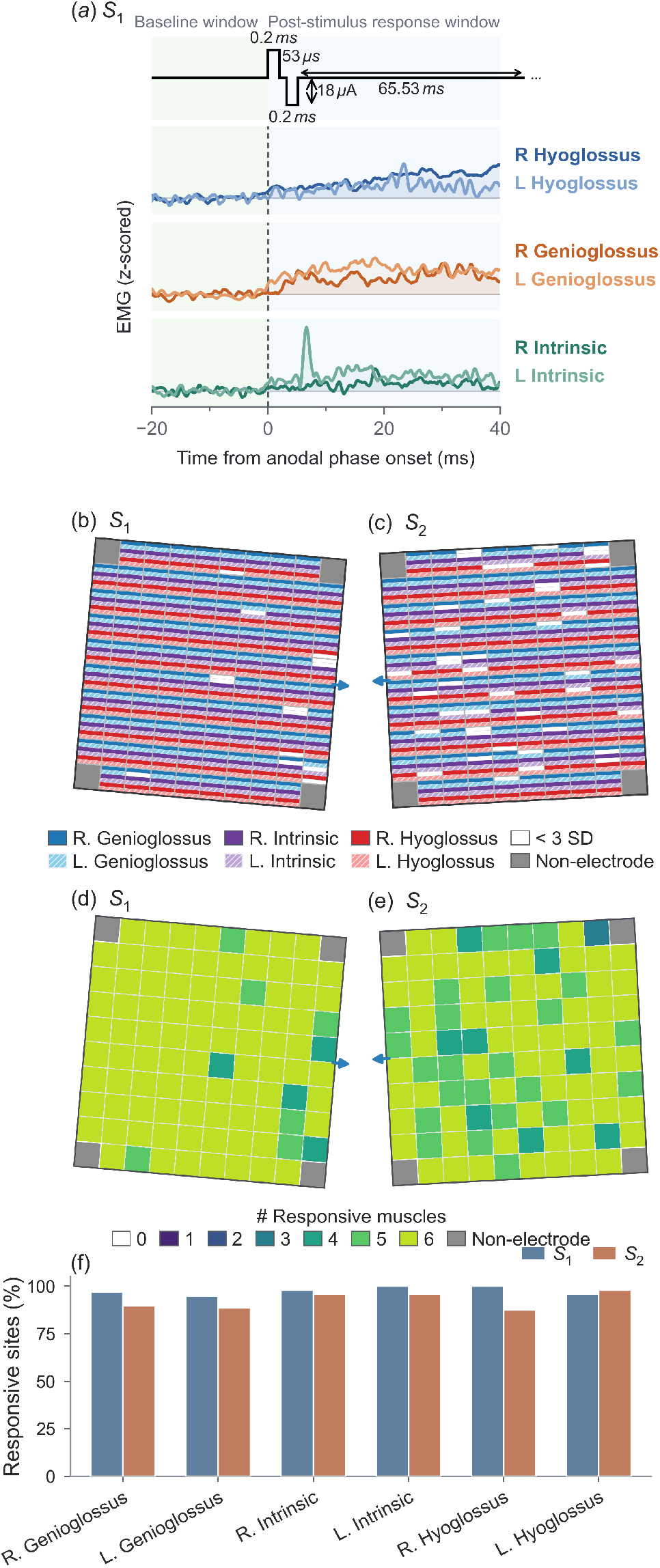
Distributed and overlapping M1 tongue-muscle output maps under intact oral sensation. (a) Representative baseline-normalized StTA responses from bilateral tongue muscles in subject *S*_1_, aligned to ICMS. Time zero denotes anodal-phase onset; traces include a 20-ms pre-stimulus baseline and 40-ms post-stimulus window. (b,c) Composite control maps for *S*_1_ and *S*_2_. Each stimulation site is divided into six bands representing, from top to bottom, right and left genioglossus, right and left intrinsic, and right and left hyoglossus. Colored bands indicate peak StTA responses ≥ 3 SD above the mean pre-stimulus EMG; white bands indicate subthreshold responses. Gray corners denote non-electrode positions, and arrows indicate the array wire-bundle direction. (d,e) Number of tongue muscles meeting the response criterion at each M1 site in *S*_1_ and *S*_2_. (f) Percentage of M1 sites evoking responses in each tongue muscle for both subjects.

Despite this widespread overlap, the overall map differences (*d*_*rms*_) and spatial distribution (r) were significant for all paired combinations of tongue muscles (see Supp Info), indicating robust representation at the level of individual muscles. Thus, intact orofacial M1 exhibited a distributed and highly overlapping tongue muscle representation, with muscle-specific differences embedded within this shared organization. These control maps provided the baseline for assessing changes produced by combined and selective oral sensory nerve blocks.

### Combined oral sensory nerve block alters ICMS-evoked tongue muscle output maps

Mechanosensory information as the tongue moves and comes in contact with oral surfaces is used to inform M1 of the sensory consequences of recently generated movement for ongoing motor control ^46–48^. For example, using the tongue to dislodge food stuck between the teeth requires continuous sensory feedback to guide the force and direction of each stroke. To test the sensorimotor loop mechanistically, we eliminated all sensory input while muscles were artificially activated using ICMS (‘Combined-NB’). In the absence of expected sensory input following the evoked muscle activity, we hypothesized a weakening of the M1-to-muscle signal and re-organization of M1 muscle maps. We quantified between-condition dissimilarity in M1-to-muscle output maps (Fig. 3a-f) using the root-mean-square electrode-wise difference, *d*_*rms*_, response magnitude, and spatial pattern similarity, *r* (see Methods).

**Figure 3.**
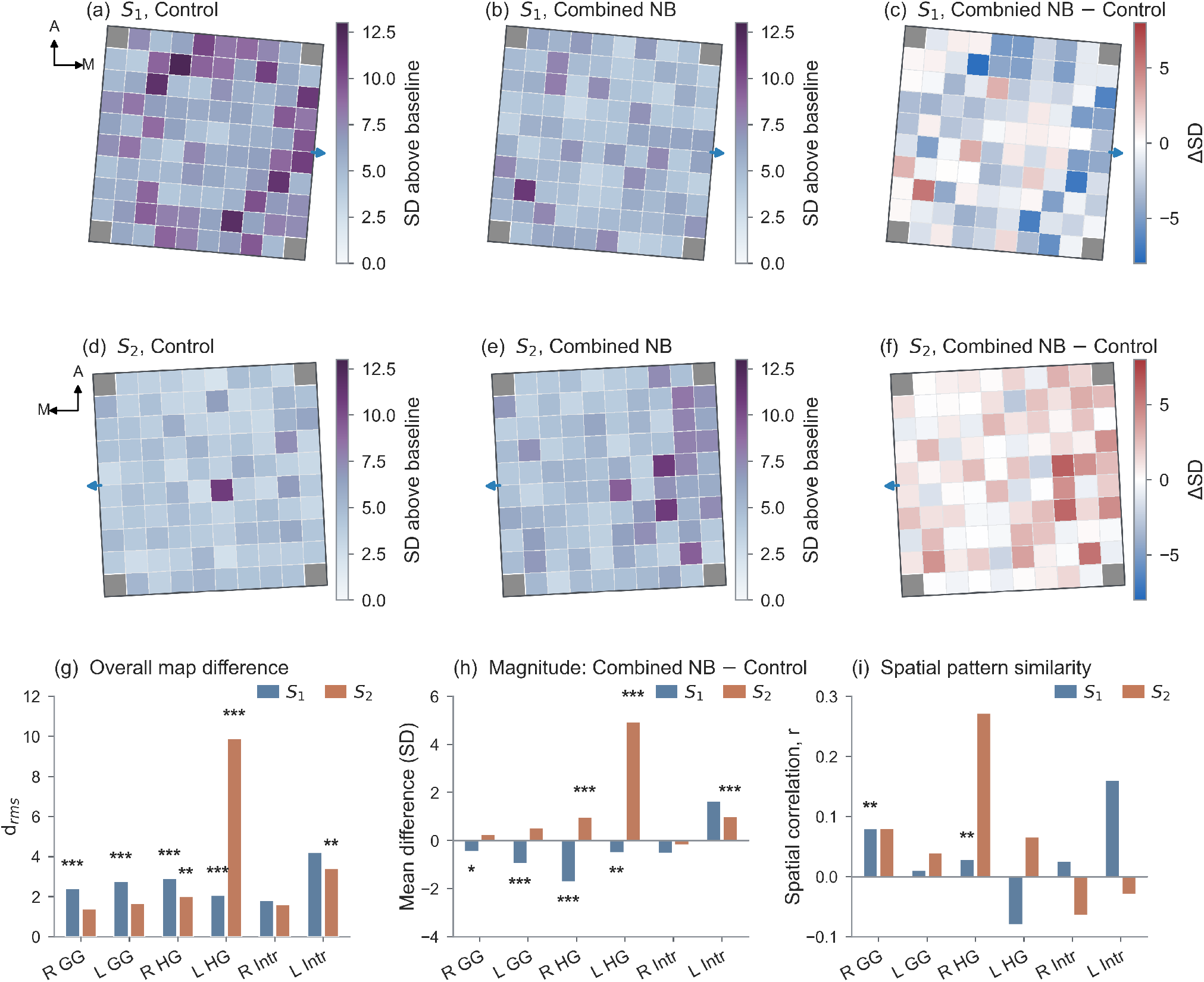
Combined oral sensory block shows muscle specific changes in tongue muscle output maps: (a-b) StTA maps of Right hyoglossus muscle for subject *S*_1_ during (a) Control and (b) Combined-NB. Map color represents the peak baseline normalized StTA response at each of 96 cortical stimulation sites, expressed as standard deviation (SD) above baseline. (c) Difference map (Combined NB - Control) for *S*_1_. Positive and negative values indicate increased and decreased responses relative to Control. (d-f) As in a-c, for subject *S*_2_. (g) Overall map difference between conditions, quantified by the root mean square difference (*d*_*rms*_). (h) Change in mean response magnitude, calculated as Combined-NB minus control; positive and negative values indicate overall enhancement and attenuation respectively. (i) Spatial pattern similarity between conditions, quantified by Pearson’s correlation coefficient across stimulation sites. Bars show the observed values for *S*_1_ and *S*_2_. Statistical significance was assessed using condition label permutation tests: right-tailed for *d*_*rms*_, two-tailed for the mean difference and left-tailed for spatial correlation. ∗ *p <* 0.05, ∗∗ *p <* 0.01, ∗ ∗ ∗ *p <* 0.001. GG, genioglossus; HG, hyoglossus, Intr, intrinsic tongue muscle.

In Combined-NB, ICMS produced significant overall map differences in at least three out of six muscles in both subjects (Fig. 3g, *d*_rms_, *p <* 0.05). These muscle-specific changes reflected shifts in the magnitude and/or spatial distribution of EMG responses (Fig. 3h-i). We illustrate these changes for the *right hyoglossus*; in subject *S*_1_, StTAs were weaker and fewer sites were responsive after nerve block (Fig. 3b) compared to the broadly distributed map under intact sensation (Fig. 3a). The resulting difference map was predominantly negative across antero-medial M1 (Fig. 3c Combined-NB − Control, right and upper-central electrodes), with localized positive values laterally. Dissimilarity measures between Control and Combined-NB were highly significant (*d*_*rms*_ = 2.9, *p <* 0.001; mean magnitude = -1.71 SD, *p <* 0.001; *r* = 0.029, *p <* 0.01), indicating both reduced response magnitude and spatial rearrangement. Similar significant changes were observed in other muscles (Fig. 3g-i, *S*_1_).

In subject *S*_2_, the right hyoglossus showed the opposite pattern: Combined-NB strengthened and broadened responses relative to the sparser control map (Fig. 3d-e), yielding a predominantly positive difference map and increased response magnitude (Fig. 3f-i, *d*_*rms*_ = 2.0, *p <* 0.01; mean magnitude = +0.98 SD, *p <* 0.001; *r* = 0.27, *p* ≥ 0.05). Several factors may explain this opposing pattern between subjects. First, baseline differences in corticomotor connectivity strength, motor unit recruitment thresholds, or resting muscle tone could set the direction of change: *S*_1_’s control map was already near a response ceiling (broad, strong baseline), leaving less room for further increases and more room for attenuation after sensory loss. In contrast, *S*_2_’s control map was comparatively sparse and weak, leaving more room for enhancement, consistent with floor effect, and potentially reflecting disinhibition of previously weak or suppressed pathways. Broadly, depending on pre-existing circuit balance, the *same* sensory manipulation may unmask facilitatory pathways in one animal and remove net excitatory drive in the other. Second, differences in the placement of cortical arrays and fine wire EMGs may contribute; because the hyoglossus fascicles follow regionally distinct fiber orientations, consistent with the muscle’s role in tongue depression and retraction ^49^, fine-wire recordings from different regions may sample distinct local patterns of muscle activity ^22,50^.

In both subjects, combined-NB affected bilateral hyoglossus response magnitude in the same direction (enhancement or attenuation) on both sides, but with greater magnitude in the contralateral hyoglossus than the ipsilateral hyoglossus (Fig. 3h), consistent with bilateral but mostly contralateral M1 projections to hypoglossal nuclei. Similar significant changes were observed across other muscles, though not uniformly: genioglossus changes were significant only in S1, and a significant intrinsic-muscle change was confined to the left intrinsic in S2, underscoring that the *direction* of reorganization is both animal- and muscle-specific.

### Combined versus selective nerve block reveals distinct muscle-specific maps

The lingual nerve carries general somatic sensation (touch, pressure, pain, and temperature) from the mucosa of the anterior two-thirds of the tongue and proprioceptive inputs from tongue muscle spindles. To determine the specific role of lingual afference in ICMS-evoked activation of tongue muscles, we blocked all sensory inputs while selectively sparing lingual inputs and compared the results against the Combined-NB condition.

When lingual afferent signals were preserved (‘Selective-NB’), a majority of the tongue muscles in both subjects exhibited significant overall map differences (Fig. 4g, *d*_rms_, *p <* 0.05). For example, the right hyoglossus motor map in Selective-NB significantly differed from Combined-NB in both subjects (Fig. 4a vs 4b and 4d vs 4e, *S*_1_:*d*_rms_ = 2.3, *p <* 0.001, *S*_2_: *d*_rms_ = 2.09, *p <* 0.05). In *S*_1_, Selective-NB increased responses at several anterolateral, central, and posterior-central sites but reduced responses at other sites. This is illustrated in Fig. 4c as the Selective-NB minus Combined-NB difference map, which contained intermixed positive and negative values, indicating spatially heterogeneous changes. In this case, the overall map difference was significant and was driven by the redistribution of responses across stimulation sites rather than by a uniform magnitude shift (Fig. 4h-i, spatial similarity: *r* = −0.22, *p <* 0.001, response magnitude: +0.28 SD, *p* ≥ 0.05). In *S*_2_, prominent Combined-NB (Fig. 4d) responses in the central-lateral and posterolateral portions of the array were reduced during Selective-NB (Fig. 4e), whereas localized increases occurred at several anteromedial and anterior sites (Fig. 4f). This was accompanied by a significant overall map difference, a modest reduction in mean response magnitude (− 0.38 SD, *p <* 0.05) and a significant reduction in spatial similarity (*r* = −0.043, *p <* 0.05).

**Figure 4.**
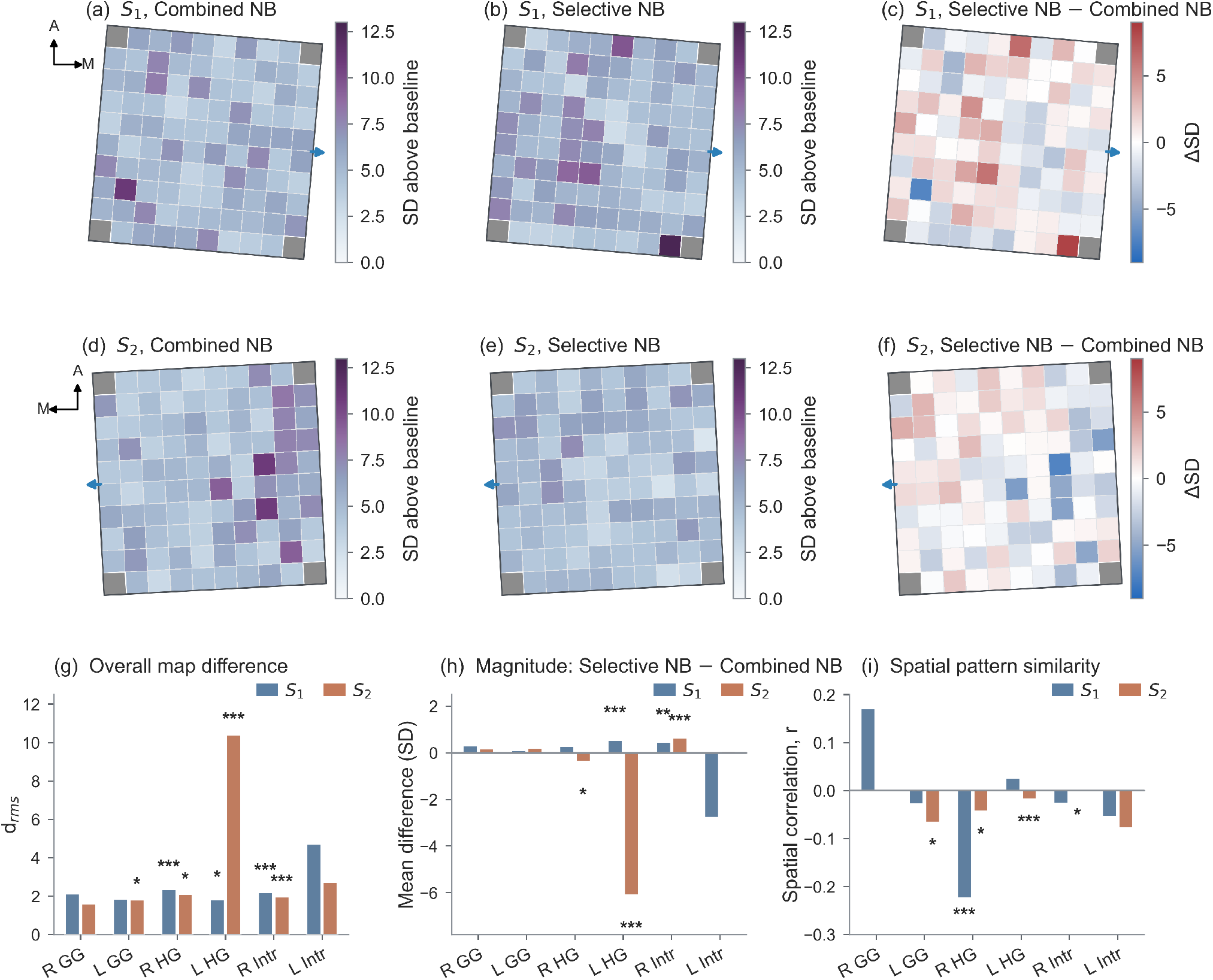
Preserved lingual afference produces muscle-specific changes in tongue-muscle output maps. **(a–c)** StTA maps of the right hyoglossus muscle for subject *S*_1_ during **(a)** Combined-NB and **(b)** Selective-NB. The corresponding difference map (Selective-NB minus Combined-NB) is shown in **(c). (d–f)** Corresponding right hyoglossus maps for subject *S*_2_. **(g)** Overall map difference between conditions, quantified by the root-mean-square difference (*d*_rms_). **(h)** Change in mean response magnitude, calculated as Selective-NB minus Combined-NB; positive and negative values indicate overall enhancement and attenuation, respectively. **(i)** Spatial-pattern similarity between conditions, quantified by Pearson’s correlation coefficient across stimulation sites. Bars show the observed values for *S*_1_ and *S*_2_. Statistical significance was assessed using condition-label permutation tests: right-tailed for *d*_rms_, two-tailed for the mean difference and left-tailed for spatial correlation. ^∗^*p <* 0.05, ^∗∗^*p <* 0.01, ^∗∗∗^*p <* 0.001. GG, genioglossus; HG, hyoglossus; Intr, intrinsic tongue muscle.

In summary, cortical output maps for the hyoglossus, genioglossus, and intrinsic tongue muscles differed significantly between the Combined-NB and Selective-NB conditions in both subjects. Notably, the hyoglossus showed overall map differences bilaterally in both subjects, suggesting a particularly robust sensitivity to lingual sensory input in this muscle’s representation. These overall map differences reflected either enhancement or attenuation of response magnitude, in some cases accompanied by spatial re-organization of the cortical-to-muscle map. Together, these findings show that lingual afference actively shapes the functional organization of tongue muscle-specific representations in M1. However, this comparison alone cannot determine whether sparing lingual afference would fully preserve M1-to-muscle output maps, or only partially supporting a graded, input-dependent model of cortical map maintenance.

### Sparing the lingual nerve differently shapes M1-to-muscle output maps

If lingual input alone were sufficient to maintain M1-to-muscle output, the cortical maps in selective-NB would not be significantly different from control. However, our results did not support this; preservation of lingual inputs still produced changes in the magnitude and spatial distribution of a majority of the tongue muscles in both subjects (Fig. 5). This indicates that loss of non-lingual oral inputs leave a distinct signature by reshaping M1-to-muscle output beyond lingual afference alone.

**Figure 5.**
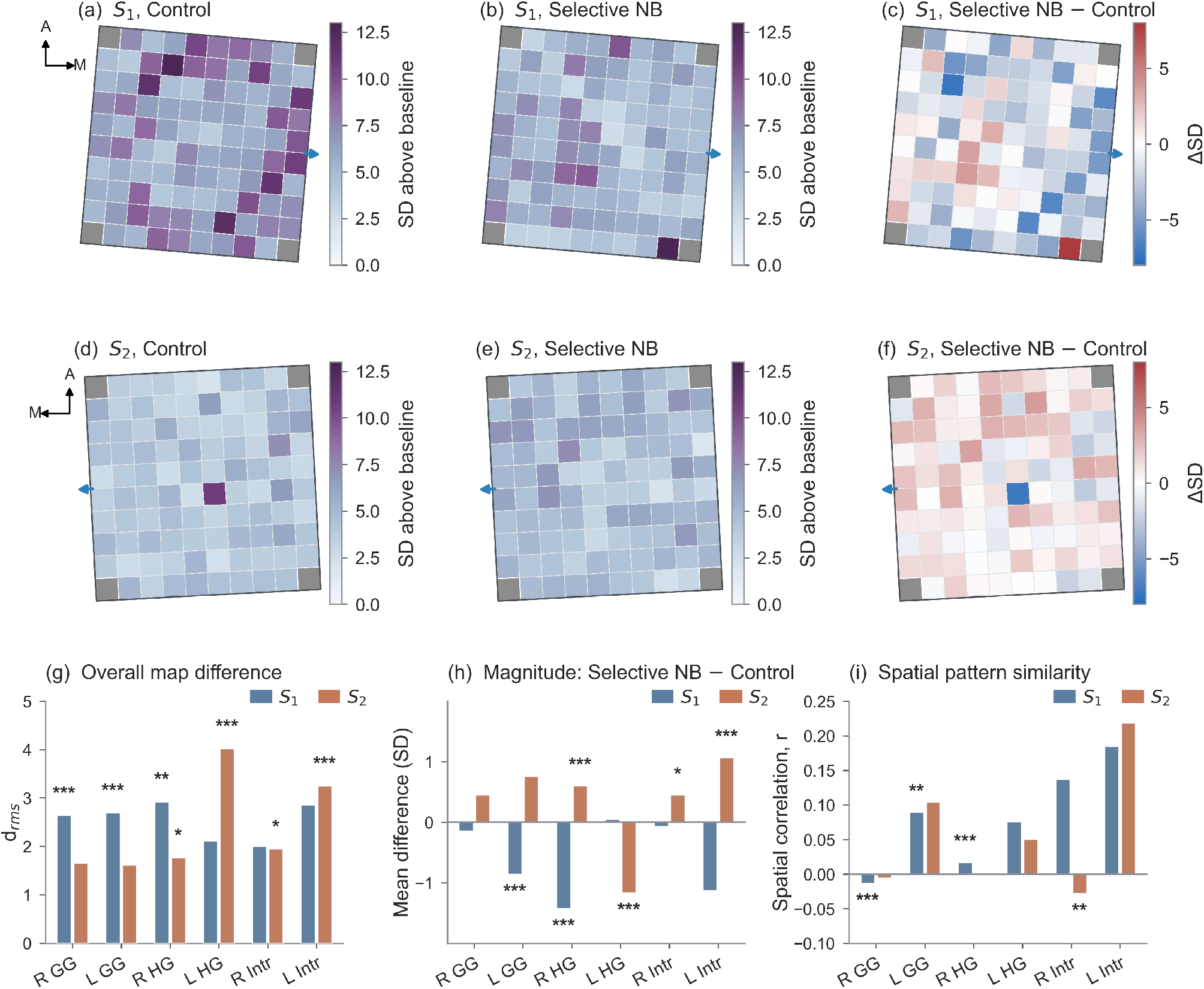
Selective oral sensory blockade produces muscle-specific changes in tongue-muscle output maps. (a–c) StTA maps of the right hyoglossus muscle for subject *S*_1_ during (a) Control and (b) Selective-NB. The corresponding difference map (Selective-NB minus Control) is shown in (c). (d–f) Corresponding right hyoglossus maps for subject *S*_2_. (g) Overall map difference between conditions, quantified by the root-mean-square difference (*d*_rms_). (h) Change in mean response magnitude, calculated as Selective-NB minus Control; positive and negative values indicate overall enhancement and attenuation, respectively. (i) Spatial-pattern similarity between conditions, quantified by Pearson’s correlation coefficient across stimulation sites. Bars show the observed values for *S*_1_ and *S*_2_. Statistical significance was assessed using condition-label permutation tests: right-tailed for *d*_rms_, two-tailed for the mean difference, and left-tailed for spatial correlation. ^∗^*p <* 0.05, ^∗∗^*p <* 0.01, ^∗∗∗^*p <* 0.001. GG, genioglossus; HG, hyoglossus; Intr, intrinsic tongue muscle.

In contrast to Combined-NB, where bilateral hyoglossus responses changed in the same direction with greater magnitude contralaterally (Fig. 3), Selective-NB produced opposing, laterality-dependent effects: in both subjects, the contralateral hyoglossus (i.e., RHG for *S*_1_, LHG for *S*_2_), showed response reduction while the ipsilateral hyoglossus showed enhancement (Fig. 5h). The contralateral attenuation of evoked responses suggests that this muscle may rely on other lingual mechanosensory inputs (floor of the mouth, lingual gingiva) for calibration while the ipsilateral enhancement suggests disinhibition by the contralateral drive. Such coupling may reflect a rebalancing or reweighting process required by symmetric activation of bilateral hyoglossus.

Across the six muscles, both subjects showed significant overall map differences for most muscles (Fig. 5g) but the underlying pattern diverged, showing either attenuation or enhancement, coupled with spatial reorganization in some cases (Fig. 5h-i). Beyond the shared effects seen in contralateral hyoglossus, the subjects differed in which muscles were affected; significant overall map differences were observed in bilateral genioglossus in subject *S*_1_ and bilateral intrinsic muscles in subject *S*_2_. Significant magnitude reductions accompanied spatial reorganization of the left genioglossus and right hyoglossus maps in subject *S*_1_, while response enhancements and minimal spatial rearrangement characterized muscle maps in subject *S*_2_.

Taken together, the findings indicate that lingual afference alone is insufficient to maintain normal M1-to-muscle output maps and that sensory inputs from other oral structures also contribute to their organization.

## Discussion

Primate orofacial M1 contains distributed and overlapping representations of the face, jaw, and tongue ^24,25^, but whether oral somatosensation modulates muscle-level response magnitude and spatial organization of tongue output from M1 has remained unknown. Here, we addressed this knowledge gap by comparing ICMS-evoked EMG responses under intact sensation and two reversible oral nerve block conditions. The resulting M1-to-muscle maps depended on which oral somatosensory input remained available; deafferentation independently altered the magnitude and spatial organization of the response at a muscle-specific level, producing qualitatively distinct output states, not graded changes. These results indicate that the muscle-level output of orofacial M1 is not a fixed property of the cortical map, but is actively and continuously shaped by the specific composition of oral sensory input available at any given time. Thus, oral somatosensation is a determinant, rather than merely a modulator, of tongue motor cortical organization.

### Distributed, many-to-many cortical organization of tongue muscles

Simultaneous mapping of bilateral genioglossus, hyoglossus, and intrinsic muscles revealed a many-to-many organization; each muscle could be recruited from numerous cortical sites and each site could recruit several muscles with different relative strengths. This organization resembles the distributed muscle-output fields identified using multi-muscle EMG mapping in macaque forelimb M1^51^, and is consistent with the broadly distributed corticospinal neurons that project to individual hand muscles ^16^. Such organization could support coordinated actions through distributed motor pathways, as suggested by stimulation studies in macaque motor cortex and by parallel corticobulbar pathways in the mouse orofacial motor system ^52,53^. It is also compatible with forelimb muscle-synergy models in which cortical stimulation recruits weighted muscle combinations ^54,55^. Nevertheless, the present data demonstrate distributed multi-muscle representation. Further study is needed to determine whether M1 representations of tongue-muscle synergies are fixed or dependent on behavioral factors.

Unlike upper limb M1, stimulation of orofacial M1 frequently evoked bilateral, multi-muscle responses, consistent with bilateral projections of M1 neurons to the hypoglossal nuclei. Multi-muscle activation may reflect direct projections from orofacial M1 neurons to the motoneuron pools of multiple tongue muscles, enabling synergistic or antagonistic co-activation; alternatively, current spread may have recruited neighboring neurons projecting to motoneurons of other muscles. While our data cannot distinguish between these mechanisms, the results support robust muscle-specific representation in M1 that is nonetheless capable of generating a coordinated tongue movement through the synchronized activity of multiple muscles.

### Oral somatosensory input shapes tongue-muscle output in a pathway- and muscle-specific manner

Temporary sensory deafferentation has previously been shown to produce divergent effects on motor-cortical output. Ischemic forearm nerve block in humans rapidly increases motor-evoked responses and can expand proximal-muscle representations ^27,28^. These effects have been attributed to disinhibition or the unmasking of pre-existing cortical pathways. In contrast, anesthetizing a defined area of skin can reduce the cortical representation of the associated muscle, while sparing or enlarging the representation of a nearby muscle ^56^. Oral deafferentation shows similarly mixed effects; lingual nerve anesthesia produces delayed facilitation of human tongue corticomotor responses ^36,37^, whereas topical tongue anesthesia yields no consistent group-level effect and substantial inter-individual variability ^57^. Our findings extend this picture of variability; attenuation, enhancement, and spatial reorganization all occurred following manipulation of sensory inputs, and that changes in response magnitude are dissociable from changes in spatial organization. Because Control mapping was performed between the two nerve block sessions rather than first or last, a simple monotonic fluctuations of array performance cannot by itself account for the pattern of results, in which Control showed the most distributed, highest-amplitude map and both flanking nerve block conditions showed attenuation or reorganization relative to it.

Therefore, oral deafferentation did not produce a uniform change in corticomotor output but instead generated muscle-specific effects, consistent with prior evidence that removing sensory feedback can either reduce muscle activity ^58^ or facilitate corticomotor output depending on context ^37^.

This muscle-specificity is consistent with, and extends, a body of pathway-specific work establishing distinct contributions of individual oral sensory territories to tongue control. Lingual afferents influence human tongue corticomotor excitability and contribute to deglutitive tongue movement ^36,37,59^ and tongue-jaw coordination during mastication in pigs ^60^. Non-lingual oral input also contributes; periodontal anesthesia alters the pressure of the tongue during swallowing ^61^, while palatal and mucosal anesthesia alters the timing of sucking-swallow, tongue kinematics, and initiation of swallowing ^62–64^. However, these studies generally measured aggregate behavioral or physiological outcomes, without resolving the effects of manipulating sensory pathways on individual tongue muscles. Our design addresses this gap directly by isolating the specific contribution of lingual afference by comparing Combined-NB and Selective-NB *within the same animals and cortical maps*, allowing pathway-specific effects to be identified at the level of individual cortical sites and individual muscles. Of special note is the consistency of the bilateral pattern of hyoglossus response across both nerve block conditions and both subjects, suggesting that hyoglossus corticomotor output is particularly sensitive to changes in sensory input, potentially reflecting its role in the precise bilateral coordination required for swallowing.

Preserving lingual afference did not uniformly restore control-like output in the present study. Selective-NB prevented some magnitude changes caused by Combined-NB but left others intact, reversed the direction of selected hyoglossus effects, and introduced spatial reorganization. These findings indicate that the anatomical composition of the remaining trigeminal afference, not merely its overall amount, determines how effectively individual cortical sites recruit particular tongue muscles. This result is difficult to reconcile with models that treat deafferentation as a graded reduction in sensory drive; if amount of input were the operative variable, Selective-NB might be expected to shift toward the Control pattern relative to Combined-NB. Instead, the Selective-NB pattern indicates that M1 output is organized around the specific identity of converging sensory channels, not simply their summed magnitude, a distinction with direct implications for how sensory loss should be modeled in motor-cortical systems. Lastly, the direction-of-change diverging between subjects for the same muscle (e.g., right hyoglossus attenuates in S1, enhances in S2) under the identical order of conditions argues against monotonic fluctuations in array performance as driving the changes.

### Candidate mechanisms and pathways

Because the nerve block effects emerged rapidly, they most likely reflect functional reweighting of existing sensorimotor pathways (e.g., via rapid changes in GABA-related inhibition or neuronal or synaptic excitability ^29^) rather than the formation of new long-range connections. These mechanisms have been characterized primarily in limb systems, but several lines of evidence suggest that comparable pathways could carry oral sensory loss to M1; for example, somatosensory manipulation alters excitability in human S1 and M1^65^, physiologically effective S1-to-M1 projections have been demonstrated in mice ^66^, coordinated interactions between orofacial S1 and M1 have been observed in macaques ^10^, and reversible inactivation of macaque face S1 also impairs tongue motor control ^33^. Direct thalamic input to M1 provides another potential route through which oral sensory loss could alter motor output ^67^. Which of these pathways drives the present effects, and to what degree, remains unresolved.

The opposing changes across animals and muscles are consistent with evidence that identical deafferentation can affect functionally distinct outputs differently. For example, ischemic nerve block increases corticospinal output to human wrist flexors but not extensors ^68^. The direction of reorganization may therefore depend on both the altered sensory pathway and the muscle examined, with additional variability potentially arising from individual differences in baseline sensorimotor state, cortical organization, or in the duration and completeness of deafferentation. Sensory loss is thus unlikely to impose a uniform increase or decrease in M1 excitability. Instead, it appears to reweight distributed sensorimotor pathways in a site- and muscle-specific manner.

### Limitations

Several limitations constrain this interpretation. First, the study included only two subjects. Second, sensory loss was short-term and reversible, making the findings most relevant to rapid functional reweighting rather than chronic deafferentation which can produce sustained reorganization of motor representations ^30–32^. Third, ICMS and stimulus-triggered EMG measure the effective output available from a cortical site rather than the output naturally recruited during behavior, thus, the relationship between these maps and muscle recruitment during feeding, swallowing, or voluntary tongue movements remains unresolved. Fourth, each experimental condition was recorded on a separate day, and the order of conditions was fixed and identical across both subjects (Combined-NB, then Control, then Selective-NB) rather than randomized or counterbalanced. Thus, the observed map differences cannot be formally dissociated from time-dependent changes unrelated to sensory manipulation. Fifth, because Control did not include sedation or reversal, residual drug effects cannot be excluded in comparisons between Control and Combine/Selective-NB. Sixth, although the tongue contains four bilateral intrinsic muscles with distinct fiber orientations and biomechanical roles, our shared cross-animal analysis included only genioglossus, hyoglossus, and one intrinsic muscle common to both subjects; findings may therefore not generalize to intrinsic muscles or extrinsic muscles not sampled here. Lastly, the study also did not record from intermediate sensorimotor pathways, so the observed changes cannot be localized exclusively to M1. Simultaneous recordings from orofacial S1, M1, thalamic nuclei, and brainstem premotor circuits will be required to identify the pathways responsible for this reorganization.

### Conclusion

These findings demonstrate that muscle-specific cortical output is not fixed but is continuously reweighted by afferent input, even in a muscular hydrostat lacking a rigid skeletal scaffold. Thus, oral somatosensory input is an active determinant of both the strength and spatial organization of tongue-muscle output from orofacial M1, rather than a peripheral factor that merely modulates a fixed motor map. Because the effects were pathway-specific rather than graded, determined by which afferents remained rather than how much input remained, this work reframes cortical motor output as a dynamically reconfigurable state that depends on the composition of converging sensory channels. This has direct relevance beyond the tongue system; it suggests that motor-cortical representations for any effector receiving convergent, multi-source sensory input may be similarly vulnerable to selective, rather than proportional, reorganization following partial sensory loss. This distinction has practical implications for understanding and treating orofacial sensorimotor dysfunction following nerve injury, dental procedures, or stroke, and for guiding sensory-based rehabilitation, neuroprosthetic or neurostimulation therapies aimed at restoring motor control.

## Methods

All procedures were approved by the University of Washington Institutional Animal Care and Use Committee and complied with the National Institutes of Health *Guide for the Care and Use of Laboratory Animals*. Reporting follows ARRIVE 2.0 guidelines.

### Subjects

Data were collected from two adult male rhesus macaques (*Macaca mulatta*); *S*_1_: 21 years old and 11.56 kg, and *S*_2_: 12 years old and 12.5 kg. Analysis were performed separately for each subject. Rhesus macaques were selected because of their similarities to humans in craniofacial anatomy, neurophysiology, orofacial motor control.

### Animal Housing and Training

Following standard quarantine, animals were acclimatized to the home-cage environment, food and water delivery systems, light-dark cycles, veterinary and research personnel, environmental enrichment, and social exposure to conspecifics. Animals were individually housed under controlled lighting, temperature, and noise conditions, with unrestricted access to water. They were fed biscuits twice daily and received supplemental foraging items, frozen treats, and fresh produce. Routine enrichment included music, television, and toys, and welfare was monitored regularly by husbandry staff.

After acclimatization, animals underwent chair training for 3-4 months before surgery. Once sufficient compliance was achieved, each animal underwent aseptic surgical implantation of a head post for head fixation during recordings. After osseointegration and bone healing were confirmed, head-fixation training was introduced alongside continued chair training.

### EMG Electrode Implantation

Bipolar electromyographic (EMG) electrodes were fabricated from medical-grade stranded stainless steel wire (Cooner Wire, Chatsworth, CA) and implanted under general anesthesia using sterile technique. Electrode pairs were tunneled subcutaneously to the submandibular region. At each recording tip, 1-2 mm of insulation was stripped, and the electrode pair was inserted into the target muscle using an 18-gauge needle. A percutaneous connector (Omnetics Connector Corporation, Minneapolis, MN), housed in a custom titanium chamber rigidly affixed to the cranium with bone screws, provided access to the differential recording channels. An additional wire wrapped around a bone screw and implanted subcutaneously served as ground.

Electrode placement was confirmed intraoperatively by electrically stimulating individual EMG channels and observing the resulting muscle-specific contractions. Stimulation consisted of single 300-ms trains delivered at 1 Hz, with each train containing 75 pulses at 250 pulses/s. Pulse duration was 0.2 ms, interphase delay was 0.01 ms, and stimulus amplitude was 5-10 V (A-M Systems, Sequim, WA).

Subject *S*_1_ was implanted with electrodes targeting seven tongue muscle channels: left and right Hyoglossus, Intrinsic, and Genioglossus, and right Styloglossus. Subject *S*_2_ was implanted with electrodes targeting six tongue muscle channels: left and right Genioglossus, Intrinsic, and Hyoglossus. The analysis reported here were restricted to the six bilateral tongue muscle channels available in both subjects: bilateral genioglossus, hyoglossus and intrinsic tongue EMG.

### Cortical Array Implantation

Under general anesthesia, each animal was chronically implanted with silicon-based Utah Electrode Arrays (Blackrock Microsystems, Salt Lake City, UT). The analyses reported here used the 96-electrode array implanted in orofacial M1, located in the left hemisphere of *S*_1_ and right hemisphere of *S*_2_. The array had 400 *µm* inter-electrode spacing and 1.0 mm electrode length.

Implantation sites were planned using co-registered CT and MRI scans in Brainsight neuronavigation software (Rogue Research, Montreal, QC), which was used to define array orientation and insertion trajectories. Arrays were implanted using a microsurgical robot (Rogue Research) following the planned trajectories. Target sites were verified intraoperatively using cortical surface landmarks and evoked responses to monopolar surface stimulation of the orofacial M1 region. Surface stimulation consisted of five consecutive 200-ms trains delivered at 5 Hz with each train containing 10 pulses delivered at 50 Hz. Pulse duration was 200 *µ*s, and stimulus amplitude was 2-4 mA.

### Experimental Design and Nerve Block Conditions

To isolate how oral somatosensory feedback influences motor cortical output, ICMS was combined with reversible peripheral sensory nerve blocks. All sessions were conducted in a dedicated acoustically controlled recording room. Subjects were seated in a customized primate chair with the head restrained to minimize movement artifacts during neural and EMG recordings.

ICMS mapping was performed under three recording sessions, one per day, with the order of conditions being fixed and identical across both subjects: Combined-nerve block, then Control, then Selective-nerve block. In the *control* condition, baseline recordings were obtained without nerve block. In the *Combined nerve block* condition, bilateral nerve blocks targeted the lingual, inferior alveolar, buccal, and maxillary nerves ^35^. In the *selective nerve block* condition, the inferior alveolar, buccal and maxillary nerves were blocked while intentionally sparing the lingual nerve. Thus, Selective-NB preserved lingual nerve afference while silencing tactile sensory input from the other targeted oral territories.

Nerve blocks were administered under sedation with ketamine (40 mg, intramuscular) and dexmedetomidine (0.15 mg for *S*_1_ and 0.1 mg for *S*_2_, intramuscular). Bilateral regional nerve block injections were performed using 0.5% bupivacaine with 1:200,000 epinephrine, with 0.25 mL administered at each injection site, following the previously described rhesus macaque oral nerve block procedure ^35^. One injection was administered for each targeted nerve on each side. Thus, Combined-NB involved eight injections (2.0 mL total), whereas Selective-NB involved six injections (1.5 mL total).

After completion of the nerve block procedure, sedation was reversed with Atipamezole (1.5 mL for *S*_1_ and 0.1 mL for *S*_2_, intramuscular), and animals were allowed to recover before ICMS mapping. Cerenia (15 mg for *S*_1_ and 12 mg for *S*_2_) was also injected subcutaneously to prevent vomiting and motion-sickness. Data acquisition was completed within 90 min of nerve block administration, within the expected period of bupivacaine efficacy ^35^. Nerve block efficacy was verified behaviorally before data collection by assessing the absence of responsiveness to tactile stimulation in the targeted oral regions. The conditions were tested in the same order in both subjects: Combined-NB, Control and Selective-NB with rest days in between recording sessions.

### ICMS and EMG Recording

ICMS was delivered using a Cerestim R96 multichannel stimulator (Blackrock Neurotech, Salt Lake City, UT) controlled by custom MATLAB scripts (MathWorks, Natick, MA). The order of electrode stimulation was randomized across the array. ICMS was delivered at 18 *µ*A and 15.15 Hz as cathodal-first, charge-balanced biphasic pulses. Each pulse consisted of a 200 *µ*s cathodal phase, a 65.535 ms interphase interval, a 200 *µ*s anodal phase, and a 53 *µ*s hardware recovery interval before the next cathodal phase. For each electrode, 34 trains of 15 pulses were delivered, yielding 510 pulses per electrode. The gap between successive trains was 1.8 ms, and the interval between stimulation of successive electrodes was 0.5 s.

During ICMS sessions, EMG activity was recorded simultaneously from implanted muscle channels. Signals were amplified and hardware-filtered using A-M Systems differential amplifiers (Sequim, WA) with a 60 Hz notch filter and a 10-500 Hz band-pass filter. EMG signals were digitized at 10 kHz.

### Stimulus-triggered EMG Analysis

Continuous EMG signals were bandpass filtered offline from 10 to 500 Hz using a fourth-order zero-phase Butterworth filter and full-wave rectified. Stimulus-triggered EMG epochs were extracted around the onset of the anodal phase of each biphasic pulse. Each epoch consisted of a 20 ms pre-trigger baseline and a 40 ms post-trigger analysis window.

Although stimulation was cathodal-first, anodal onset was used as a temporal reference for alignment. Because the anodal phase lasted 200 *µs* and was followed by a 53 *µs* recovery interval, the next cathodal phase began 253 *µs* after the trigger. The 20 ms pre-trigger baseline therefore occurred within the long interval following the preceding cathodal phase and was verified to contain no detectable stimulus-locked modulation attributable to that preceding stimulation. Each 15 pulse train provided 14 valid anodal aligned trigger events. The initial cathodal phase in each train did not have a preceding anodal trigger, whereas the final anodal onset was excluded because the complete post-trigger analysis window was not available within the stimulation train. Across 34 trains, this yielded 476 valid trigger events per electrode.

For each electrode-muscle pair, the 476 valid rectified EMG epochs were averaged sample by sample to obtain one stimulus-triggered average (StTA) waveform. The baseline mean (*µ*_*baseline*_) and standard deviation (*σ*_*baseline*_) were calculated from the 20 ms pre-trigger portion of the averaged waveform. Response magnitude was defined as the peak value in the 40 ms post-trigger analysis window, expressed relative to the baseline as

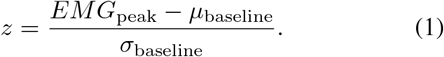

Here, *EMG*_peak_ is the peak amplitude of the averaged and rectified EMG waveform during the post-trigger window. The resulting value therefore represents the peak baseline normalized StTA response at that cortical site.

For Fig.2b-c, a cortical site was classified as responsive for a given muscle when its peak StTA response was at least 3 baseline standard deviations above the pre-stimulus mean. This threshold was used to visualize the prevalence and overlap of tongue-muscle responses across the array. The continuous StTA response magnitudes were retained for subsequent map comparisons.

Computational analysis was performed in MATLAB 2022b (MathWorks) and Python.

### Condition-wise Map Comparison

Let *A*_*e*_ and *B*_*e*_ denote the magnitude at electrode *e* under conditions *A* and *B*, respectively. Overall map dissimilarity was quantified using the root-mean-square electrode-wise difference:

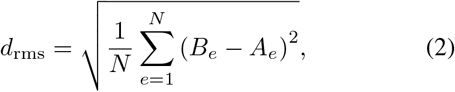

where *N* is the number of electrodes. A value of *d*_rms_ = 0 indicates identical maps, whereas larger values indicate greater overall map dissimilarity. Because *d*_*rms*_ is unsigned, it does not distinguish whether map differences arise from an overall change in response magnitude, a spatial reorganization across electrodes, or a combination of both.

Statistical significance of *d*_rms_ was assessed using a condition-label permutation test. For each muscle and electrode, the stimulus-triggered EMG epochs from the two conditions were pooled and randomly reassigned to two surrogate condition groups. The reassigned waveforms were averaged within each surrogate condition, and a new baseline-normalized peak response was recalculated for each averaged waveform per electrode. *d*_rms_ was then recalculated between these maps. This procedure was repeated 10,000 times to generate a null distribution representing map differences expected after removing original condition labels. The right-tailed permutation p-value was computed as

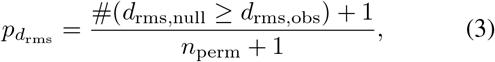

where *d*_rms,obs_ is the observed map difference, *d*_rms,null_ is the value obtained from each permutation, and *n*_perm_ = 10,000.

To determine whether map differences included an overall change in response magnitude, the signed difference in mean response was computed as:

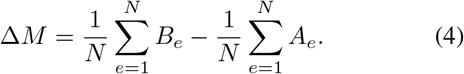

Positive values indicate larger average responses in condition *B*, whereas negative values indicate larger average responses in condition *A*. The same condition-label permutations were used to generate a null distribution for Δ*M*. Because the changes could occur in either direction, significance was assessed using a two-tailed test:

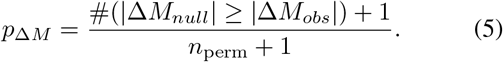

Spatial pattern similarity was quantified using the Pearson’s correlation coefficient between the two electrode-wise maps,

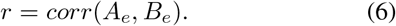

Values of *r* approaching 1 indicate that electrodes with relatively strong responses in one condition also tended to have relatively strong responses in the other condition. Because Pearson’s correlation is computed after mean-centering each map, it primarily characterizes the relative spatial pattern rather than differences in overall response magnitude. Statistical significance was assessed using a left-tailed permutation test asking whether the observed spatial correlation was lower than expected under the condition-label null distribution:

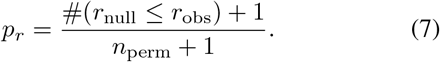

To further characterize the source of overall map differences, the squared map difference was decomposed into global and heterogeneous components. Defining the electrode-wise condition difference as

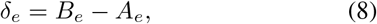

the squared RMS difference can be written as:

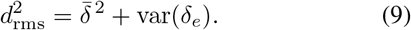

The first term, 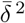, represents the magnitude contribution of the global mean shift between conditions. The second term, var(*δ*_*e*_), represents electrode-to-electrode heterogeneity in the condition effect. The fractional contribution of each component was calculated by dividing that component by 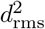.

Permutation tests were performed independently for each muscle and statistic. Resulting p-values were adjusted using the Benjamini–Hochberg procedure across the six muscles, separately for each subject, condition comparison, and statistic. Significance was defined as adjusted *p <* 0.05. All p-values reported in the Results, figures, and tables are adjusted values.

A significant *d*_rms_ indicated an overall difference between condition-specific maps. A significant Δ*M* indicated an overall increase or decrease in response magnitude, whereas a significant reduction in spatial *r* indicated lower spatial similarity between maps. These measures were interpreted together rather than as mutually exclusive descriptions of the condition effect.

### Inter-muscle Map Comparison

To determine whether individual tongue muscles had distinct cortical output maps under the Control condition, all 15 pairwise comparisons among the six bilateral tongue muscles were performed separately for each subject. The same three aspects of map organization used for the condition-wise comparisons were considered: overall map difference (*d*_rms_), global magnitude difference (Δ*M*), and spatial similarity (Pearson’s *r*), as defined above.

The condition-label permutation used for across-condition comparisons was not appropriate for inter-muscle comparisons because each EMG response must remain associated with the muscle from which it was recorded. Ordinary 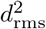 is positively biased ^69^ by measurement noise because random electrode-wise differences are squared and therefore increase the estimated distance even when the underlying maps are identical. We therefore used a cross-validated squared Euclidean distance ^70^ to determine whether the difference between two muscle maps was reproducible across separate subsets of the data.

For each electrode, all 476 valid pulse-triggered EMG waveforms were randomly divided without replacement into two non-overlapping subsets, A and B, using identical pulse assignments across simultaneously recorded muscles and a separate StTA map was constructed from each subset. For a pair of muscles, the electrode-wise difference maps were calculated as

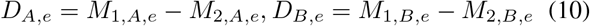

where *M*_1,*A,e*_ and *M*_2,*A,e*_ are the responses of the two muscles at electrode *e* estimated from subset A, and equivalently for subset B. The cross-validated squared distance was then calculated as

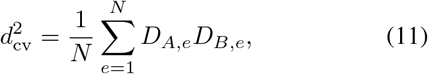

where *N* = 96 is the number of stimulation electrodes. The hypotheses is that if two muscle maps genuinely differ, the electrode-wise difference pattern should reproduce across the two subsets, producing a positive 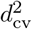. Random measurement noise should not reproduce consistently across the subsets, so under the null hypothesis of no reproducible map difference the expected 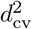 is zero. To reduce dependence on any particular random split, the split-half procedure was repeated 100 times and the resulting 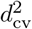 values were averaged.

Although cross-validation reduces the positive noise bias in the estimated map difference, it does not quantify the sampling variability arising from the finite number of pulse-triggered responses. We therefore used non-parametric bootstrapping to estimate this variability. For each electrode-muscle combination, all the pulse-triggered waveforms were resampled with replacement to generate a bootstrap dataset of the same size as the original dataset. The same split-half procedure was then applied, with 100 random splits averaged to obtain one 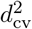 value for each bootstrap replicate. This procedure was repeated 10,000 times. A one-sided bootstrap *p*-value was calculated as

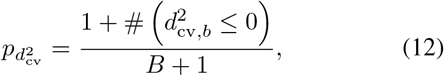

where, the bootstrap replicates, B=10,000.

The magnitude difference between the two muscles was assessed using the same 10,000 bootstrap datasets but did not require split-half cross-validation. For each bootstrap replicate, complete StTA maps were reconstructed and Δ*M* was recalculated. Because either muscle could have the greater mean response, significance was assessed using a two-sided percentile-bootstrap test against the null value of zero using the following equation ^71^,

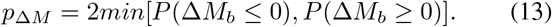

Pearson’s *r*, calculated from the complete maps, was reported as a descriptive measure of spatial similarity. Statistical evidence for differences in spatial organization was assessed separately using the cross-validated distance after removing the global magnitude component. For each split-half map, the mean response across electrodes was subtracted before calculating the muscle-difference maps. The resulting mean-centered 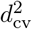 was evaluated using the same bootstrap procedure described above, with

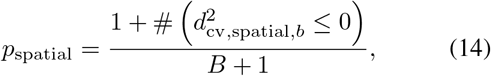

Thus, Pearson’s *r* described the similarity of the spatial patterns, whereas the mean-centered cross-validated distance provided the corresponding statistical test. For each subject and statistic, p-values were adjusted across the 15 muscle-pair comparisons using the Benjamini–Hochberg procedure, with significance defined as adjusted *p <* 0.05.

## Author contributions

F.I.A.-M. conceived and supervised the project and developed the methodology. S.P. developed the software and performed the primary data analysis and visualization. S.P., E.D.T., and F.I.A.-M. contributed to formal analysis and interpretation of the results. S.P., L.Q.F., S.H., J.S.L., and D.T. performed the experiments. A.B.B. performed the surgical procedures. H.C. performed the CBCT imaging. S.P. wrote the original draft of the manuscript. F.I.A.-M., S.P., and E.D.T. reviewed and edited the manuscript. F.I.A.-M. administered the project and acquired funding. All authors discussed the results, revised the manuscript, and approved the final version.

## Acknowledgements

We are grateful to the veterinary staff, husbandry team, surgical team, and Behavioral Management Services of the Washington National Biomedical Research Center (WaNBRC) for their dedicated support and care of the animals. We also thank Prof. Andrew Nalley from the Department of Oral and Maxillofacial Radiology at the University of Washington for his assistance with CBCT imaging.

## Data availability

The raw neural and EMG data are available from the corresponding author upon reasonable request.

## Supplementary Information

**Supplementary Fig. 1.**
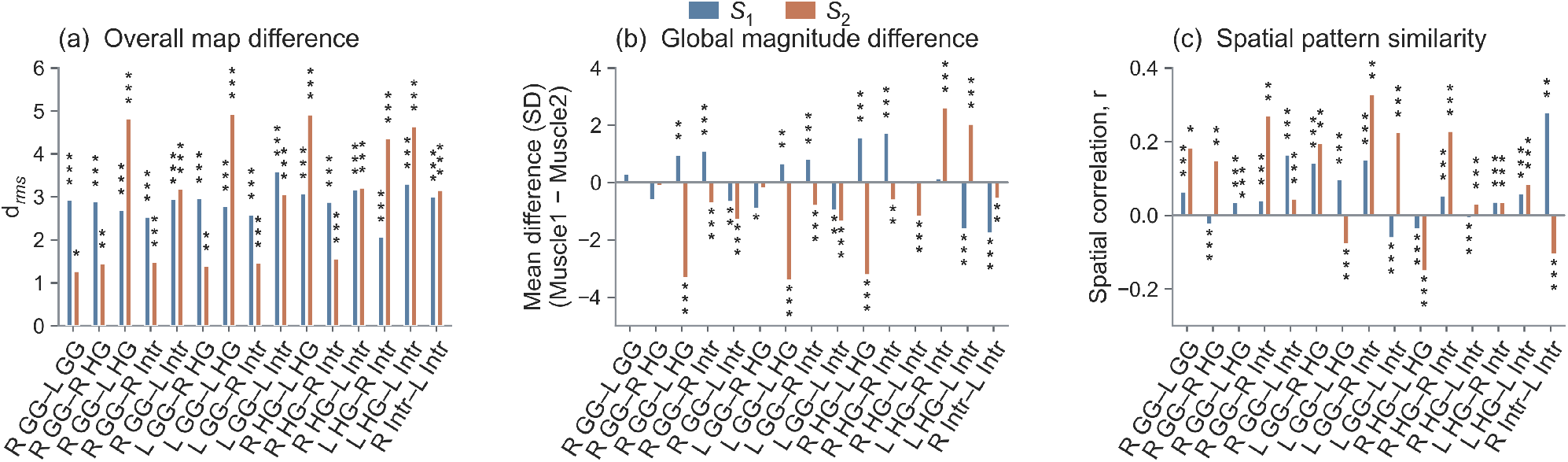
Intermuscle differences in cortical tongue-muscle output maps during the Control condition. Pairwise comparisons among the six tongue muscles are shown for subjects *S*_1_ and *S*_2_. (a) Overall map difference quantified by *d*_rms_. (b) Difference in mean response magnitude between the two muscles in each pair (Muscle 1 − Muscle 2), expressed in SD above baseline. Positive values indicate a larger mean response for Muscle 1, whereas negative values indicate a larger mean response for Muscle 2. (c) Spatial similarity between the two cortical output maps, quantified by the Pearson correlation coefficient *r*. Blue and orange bars denote *S*_1_ and *S*_2_, respectively. Significance symbols indicate FDR-corrected pairwise tests: \**p <* 0.05, \*\**p <* 0.01, and \*\*\**p <* 0.001. R, right; L, left; GG, genioglossus; HG, hyoglossus; Intr, intrinsic.

**Supplementary Fig. 2.**
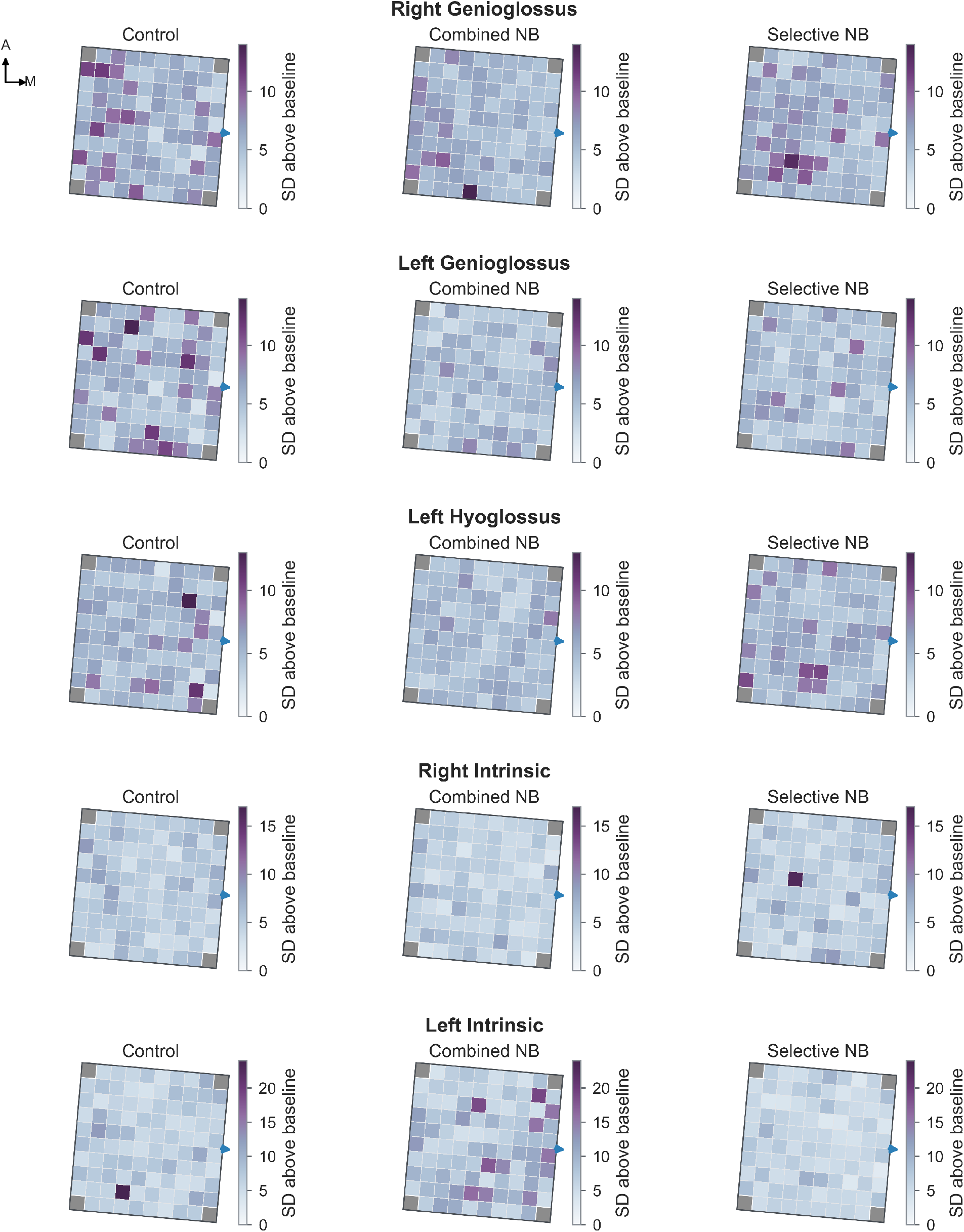
Tongue-muscle output maps across oral sensory conditions in subject. *S*_1_. StTA maps of remaining five tongue muscles: right genioglossus, left genioglossus, left hyoglossus, right intrinsic, and left intrinsic. Columns show responses during Control, Combined nerve block (Combined-NB), and Selective nerve block (Selective-NB) conditions. Color scales are held constant across conditions within each muscle to permit direct comparison. A, anterior; M, medial.

**Supplementary Fig. 3.**
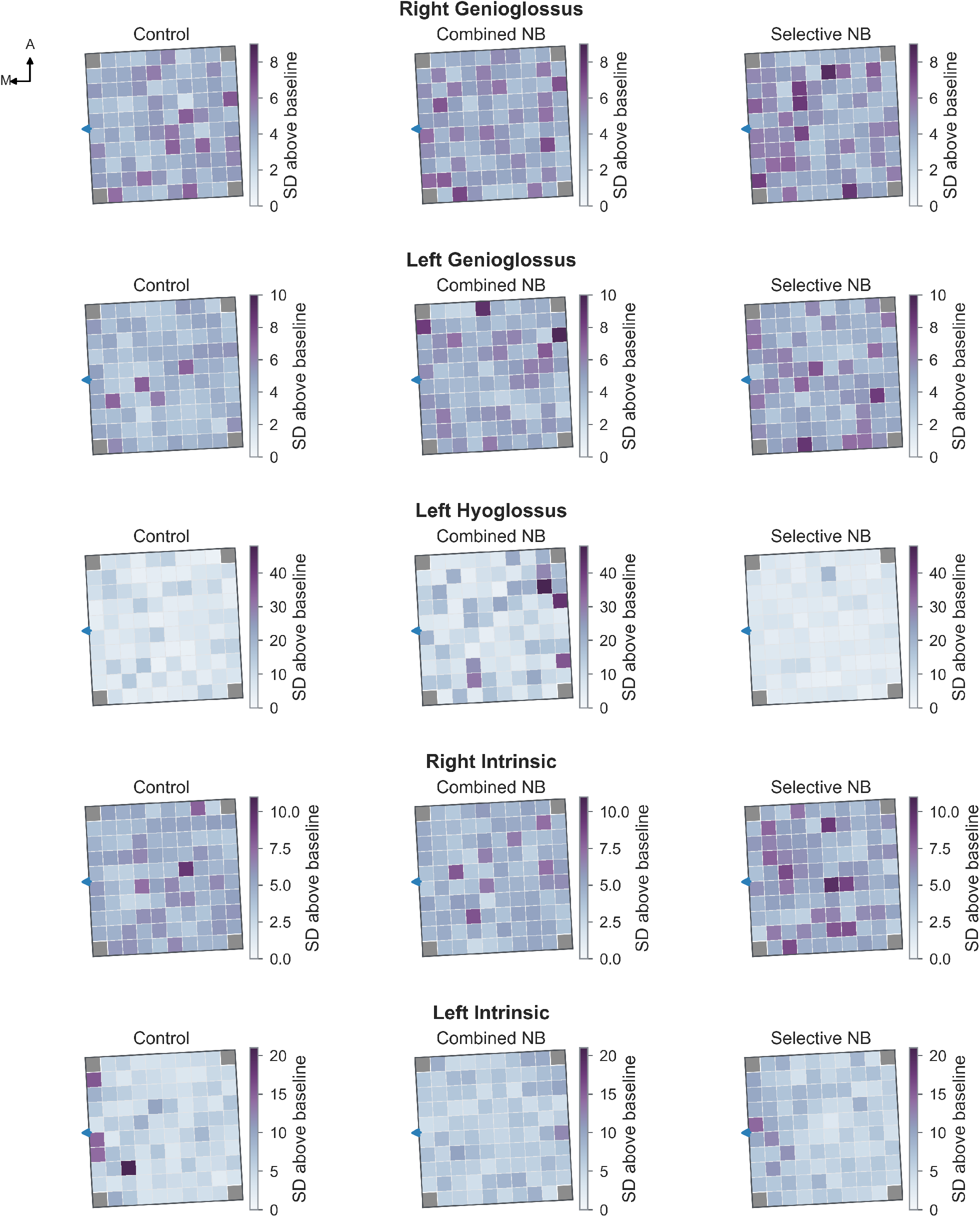
Tongue-muscle output maps across oral sensory conditions in subject. *S*_2_. StTA maps of remaining five tongue muscles: right genioglossus, left genioglossus, left hyoglossus, right intrinsic, and left intrinsic. Columns show responses during Control, Combined nerve block (Combined-NB), and Selective nerve block (Selective-NB) conditions. Color scales are held constant across conditions within each muscle to permit direct comparison. A, anterior; M, medial.

